# Dominant carnivore loss benefits native avian and invasive mammalian scavengers

**DOI:** 10.1101/2021.09.06.459188

**Authors:** Matthew W. Fielding, Calum X. Cunningham, Jessie C. Buettel, Dejan Stojanovic, Menna E. Jones, Barry W. Brook

## Abstract

Scavenging by large carnivores is integral for ecosystem functioning by limiting the build-up of carrion and facilitating widespread energy flows. However, top carnivores have declined across the world, triggering trophic shifts within ecosystems. In this study, we use a natural ‘removal experiment’ of disease-driven decline and island extirpation of native mammalian (marsupial) carnivores to investigate top-down control on utilisation of experimentally placed carcasses by two mesoscavengers – the invasive feral cat and native forest raven. Ravens were the main beneficiary of carnivore loss, scavenging for five times longer in the absence of native mammalian carnivores. Cats scavenged on almost half of all carcasses in the region without dominant native carnivores. This was eight times more than in areas where other carnivores were at high densities. In the absence of native mammalian carnivores, all carcasses persisted in the environment for 3 weeks. Our results reveal the efficiency of carrion consumption by mammalian scavengers. These services are not readily replaced by less-efficient facultative scavengers. This demonstrates the significance of global carnivore conservation and supports novel management approaches, such as rewilding in areas where the natural suite of carnivores is missing.

## Introduction

Scavenging is ubiquitous among mammalian and avian carnivores, with most species scavenging to some degree [1-4]. Larger carnivorous mammals are highly efficient scavengers, consuming carcasses faster than most other taxa [5]. However, many larger mammalian carnivores, as well as obligate scavengers like vultures, are experiencing widespread declines due to habitat loss, disturbance, and persecution by humans [6, 7]. Fluctuations in the abundance of these species can have trophic consequences that cascade throughout the food web and impact nutrient cycling and disease transmission [8-10]. With populations of some larger mammalian carnivores now beginning to recover, this raises questions around how scavenging dynamics have shifted within modified ecosystems [11].

Larger mammalian carnivores can either provision ecosystems with a more stable supply of carrion (e.g. wolves in Yellowstone National Park [12]), or limit carrion access (e.g. bears kleptoparasiting cougar kills [13] and Tasmanian devils reducing carrion availability [14]). Carrion is a high-quality resource with low handling costs, and thus is attractive to mesoscavengers [1]. However, scavenging on carrion is also risky due to the increased likelihood of encountering dominant scavengers [15-17]. These competitive and facilitative processes can potentially make carrion “fatally attractive” for mesoscavengers [18]. For example, mesoscavengers were attracted to wolf kills yet were negatively associated with wolf density at the landscape scale [19]. Although carcasses are attractive to mesoscavengers, predator avoidance plays an important role in shaping carnivore communities [18, 20].

Across the southern-temperate continental island of Tasmania (Australia) and its large offshore islands (Fig. 1; total area: 68,401 km), a large-scale natural experiment is occurring due to the severe population decline of the largest extant terrestrial carnivore, the marsupial Tasmanian devil *Sarcophilus harrisii* [20]. Devils are Tasmania’s dominant scavenger, being both the largest extant terrestrial mammalian carnivore and a specialist, although facultative, scavenger adapted for processing the toughest parts of carcasses [21]. Following the extinction of the thylacine *Thylacinus cynocephalus* in the 20^th^ century, this mesocarnivore has become the apex terrestrial carnivore [20, 22, 23]. Devils have experienced severe population declines due to a transmissible cancer, devil facial tumour disease (DFTD) [24]. The disease has progressively spread across Tasmania over 25 years, causing average population declines of 83% across ∼90% of Tasmania [25, 26]. The progressive spread of DFTD has established a natural experiment by creating regions of Tasmania with different disease histories and consequently, widely variable densities of top carnivores. Unlike most threatened carnivores [6], devil population declines are not caused by humans, allowing us to study the effects of a carnivore’s abundance with little anthropogenic confounding. In areas where devils have declined, carrion persists three-fold longer, allowing increased carrion consumption by native (spotted-tailed quolls *Dasyurus maculatus*) and invasive (feral cats *Felis catus*) mammalian and avian (forest ravens *Corvus tasmanicus*) mesoscavengers [14]. However, this prompts the question: what would happen to carrion if all native mammalian carnivores were completely lost? Can invasive and avian mesoscavengers then fully replace the ecosystem services of larger mammalian scavengers?

**Figure 1.**
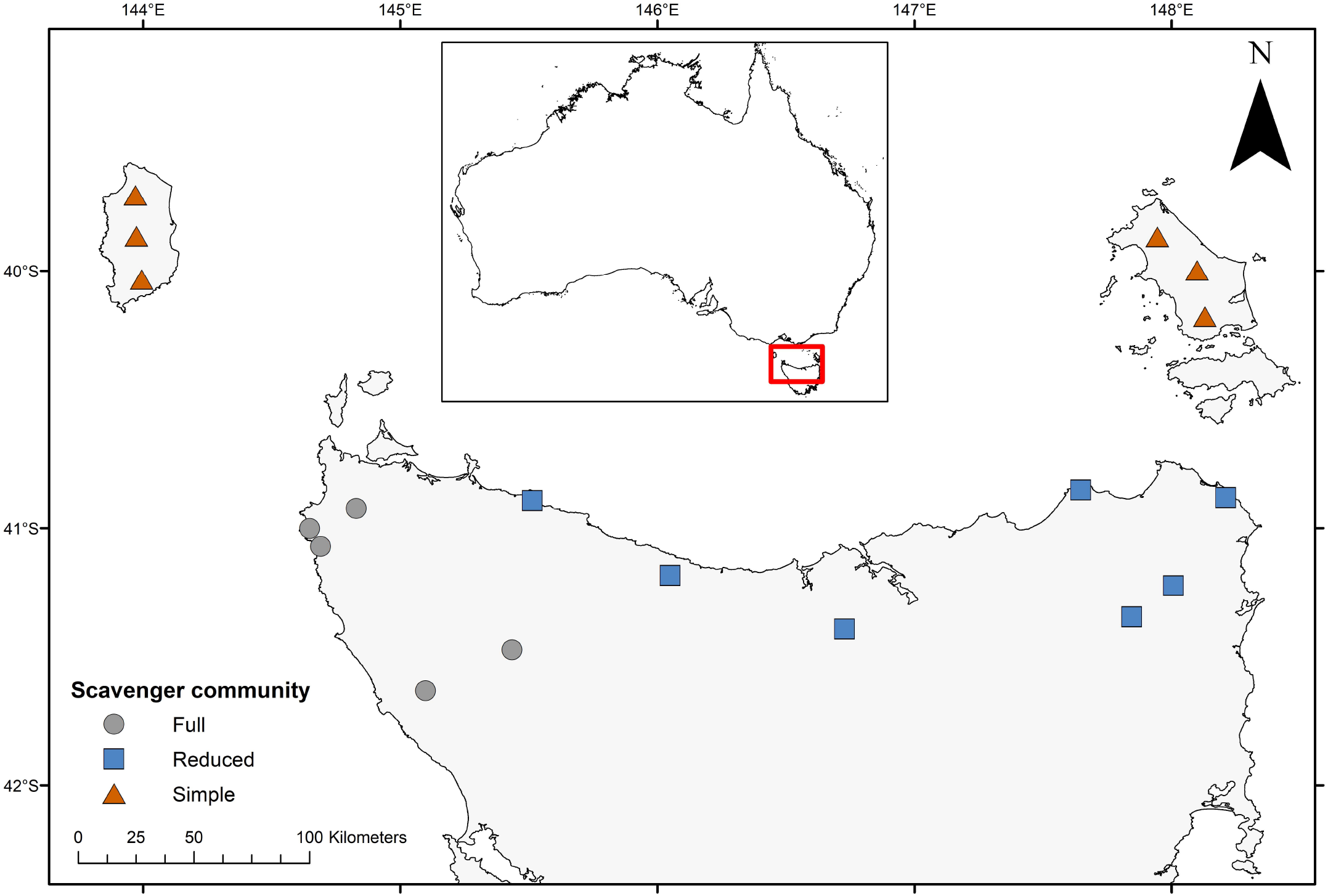
Geographic location of study sites across Northern Tasmania and the Bass Strait Islands. Each shape indicates a site that contains between 6 to 8 camera traps.

The Bass Strait Islands, between Tasmania and the Australian mainland, are ecologically similar to mainland Tasmania due to intermittent land connectedness during glacial maxima [27]. However, following major land-use change and human persecution, several species were driven to extinction on the islands, including mammalian carnivores like spotted-tailed quolls [28, 29]. While there is no evidence of the Tasmanian devil on the islands following European occupation, on Flinders Island, fossil evidence suggest that devils may have persisted until at least 8000 years ago [29, 30]. This mensurative experiment on the Bass Strait Islands thereby provides a unique opportunity to compare scavenging between: 1) a full community (Tasmanian mainland) comprising a native mammalian apex scavenger, native mammalian mesoscavenger, invasive mammalian mesoscavenger and native avian mesoscavenger; 2) a community in which only the native apex scavenger has declined (Tasmanian mainland in diseased areas); and 3) a community lacking all native mammalian scavengers (Bass Strait Islands).

In this study, we used experimentally deployed carcasses (n = 136) and camera traps to monitor carrion use by mammalian and avian scavengers across mainland Tasmania and the two largest Bass Strait Islands. Using this design, we investigated how the abundance and occurrence of native top scavengers (devils) and native mesoscavengers (quolls) impacts: (i) carrion use and discovery by invasive and extant native mesoscavengers, and (ii) carcass persistence within an environment. We hypothesised that avian generalist mesoscavengers (ravens) and invasive mesoscavengers (feral cats) will not match the scavenging efficiency of devils and predicted that carcasses should persist for longer in their absence. We also expected that top native mammalian scavengers (devils) would limit carrion access and total feeding time for other smaller scavengers. Furthermore, we hypothesised that devils would have a stronger impact on the scavenging community than smaller native mesoscavengers (quolls).

## Methods

### Study area

We measured carrion use by mammalian and avian scavengers across Tasmania and the two largest Bass Strait Islands in south-eastern Australia (Fig. 1). In Tasmania, the progressive westward spread of DFTD from its origin in the northeast, followed by rapid and severe local population decline has resulted in low devil densities across most of the state. We divided northern Tasmania and the Bass Strait islands into three regions, partitioned geographically based on the density of native mammalian scavengers: 1) a full community, where DFTD was absent or only recently invaded, and devils were abundant and quolls present although naturally at lower densities than devils; 2) a reduced community, where DFTD was prevalent and devil numbers declined by more than 80%, however, quoll densities do not appear to have increased substantially; and 3) a simple community, where devils and quolls are locally extinct (Fig. 1). In total, we experimentally deployed 136 carcasses and cameras across these regions (full community: 40; reduced community: 56; simple community: 40). Camera sites were placed in a roughly even mixture of wet eucalypt/rainforest habitat and dry eucalypt/coastal scrub habitat and we selected areas where human influence was minimal.

### Experimental design

Carcasses were deployed during August - September 2016 for the Tasmanian mainland, and August - September 2020 on the Bass Strait islands. We worked in late winter, when consumption by invertebrate scavengers and microbial decomposers is at its lowest. We used Bennett’s wallaby (*Macropus rufogriseus*) and Tasmanian pademelon (*Thylogale billardierii*) carcasses. These are both regularly culled under crop-protection permits and are a common source of carrion in the study region. The carcass species used at a given site depended on local availability. To ensure independence, carcasses were deployed at least 1 km apart. At each carcass, we installed one camera trap (Cuddeback X-Change 1279 or Reconyx PC-800) to monitor scavengers. Cameras were deployed for a minimum of 21 days, after which we expected the carcasses to be mostly consumed.

### Analysis

#### i) Carcass discovery & persistence

We used statistical survival analysis based on a mixed-effects Cox proportional hazards model from the R package `‘coxme’` [31] to study carcass discovery and persistence. We ran separate analyses to investigate the time it took for the carcasses to be discovered by (i) any vertebrate scavenger, (ii) ravens, and (iii) feral cats. Discovery was defined as the first time an animal found and fed on the carcass. Carcasses were defined as fully consumed when there was a clear final consumption event, and the physical carcass was absent from subsequent images. If the final event was not clear, we used the 95^th^ percentile foraging event, when foraging activity petered out over many days, given the carcasses appeared to be more than 95% consumed.

We used survival analysis because the discovery and persistence data were censored [32]. Discovery data were right-censored because not every carcass was discovered by ravens and feral cats before the carcass was completely consumed. The persistence data were also right-censored because memory cards occasionally became full before complete consumption (n = 9), several carcasses were prematurely removed from the view of the camera (n = 18), one camera returned no images throughout the study, and the batteries in one camera failed before the carcass was fully consumed.

We selected the preferred survival models using the package `‘MuMIn’`[33] with combinations of four predictor variables: devil activity (number of devil detections per 100 camera nights), quoll activity (number quoll detections per 100 camera nights) and habitat (wet vs dry forest), with initial carcass weight (kg) included as a covariate to account for variation in carcass size (see Table S1 in the Supplementary Information for model combinations). To account for variation across the study sites, we used site location as a random effect. All the predictors had a Pearson’s cross-correlation coefficient *r* < 0.7. Model selection was based on Akaike’s information criterion corrected for a small sample size (AIC_c_). The most parsimonious models were selected using ΔAIC_c_ < 2 (difference between the AIC_c_ of a given model and the best model) (Grueber et al. 2011). For each supported covariate (where 95% confidence limit did not overlap with zero), we calculated the model-averaged exponentiated coefficients, known as hazard ratios (HR), which provide effect sizes for each variable. Survival curves were visualised by separating carcass data into the three regions (Fig. 1) and presenting the Kaplan-Meier estimates of the survival function using the packages `‘survival’` [34] and `‘survminer’` [35].

#### ii) Carcass use

To analyse the predictors of carcass use by forests ravens and feral cats, we tested a range of *a priori* models based on ecological knowledge (Table S2, supplementary information). We again used AIC_c_ for model selection and multi-model inference. We assessed the fit of the top models by calculating the AUC (area under the receiver operator curve; suitable for classification models). We tested the effects of five predictor variables: habitat, devil, and quoll activity (defined above), plus total scavenging time by devils and quolls (separately, summated minutes). To account for variation in carcass size, we included initial carcass weight as a covariate. We also used site location as a random effect to account for variation across the study sites. We calculated the standardised regression coefficient (Std. coef) based on z-values for each predictor after model averaging. For any variables that were supported (95% confidence limit did not overlap with zero), we calculated the effect size (ES) by comparing the model-averaged predicted probability when that categorical variable was applied, against the probability of the intercept when the effect was absent. Sixteen cameras were removed from the analysis in total: twelve due to premature removal of the carcass from the field of view and four due to mechanical unreliability or early failure.

To analyse carcass use by forest ravens, we used hurdle models, because the scavenging data were zero-inflated and followed a gamma distribution. We first modelled whether ravens fed at a carcass (GLMs with binomial link function) and then modelled the total scavenging time by ravens for the carcasses at which they fed (GLMs with a Gamma distribution and a log link function). Total scavenging time for each camera was calculated by summing the number of minutes any raven spent scavenging on the carcass. GLMs with a binomial link function were used to assess carcass use by feral cats, but we were unable to model the predictors of total scavenging time by feral cats due to insufficient data, particularly in the full community region. Carcass use was defined as whether a feral cat found and scavenged upon a carcass during the study period.

## Results

### i) Carcass discovery & persistence

The absence of native mammalian carnivores had a significant effect on extending the persistence time of carcasses. Carcasses within the simple community region lasted 1.8 times longer than reduced community regions and 4.6 times longer than full community region (Fig. 2). ‘Devil activity’ had a negative effect on carcass persistence (Hazard Ratio, HR = 1.09) while ‘carcass weight’ had a positive effect, with larger carcasses persisting for longer (HR = 0.96).

**Figure 2.**
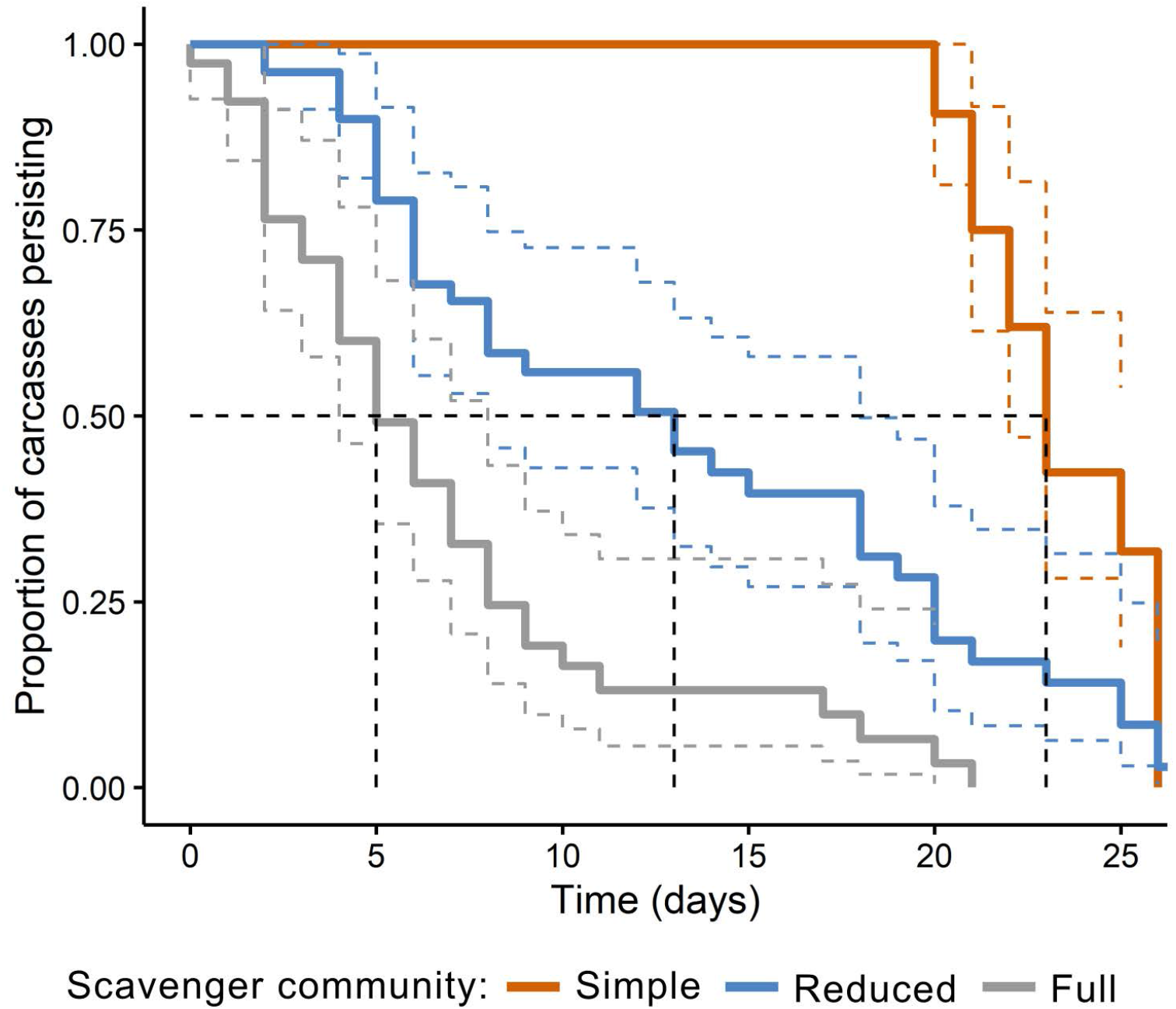
The proportion of carcasses persisting in the environment for each level of scavenger diversity and abundance. Colour dashed lines indicate the 95% confidence interval. Black dotted line shows the median carcass persistence times for each of the three categories.

There was no difference in the discovery rates between the various scavenger communities when all species were aggregated (Fig. 3a), although carcasses in wet forests took longer to be discovered (HR = 0.54). Ravens discovered carcasses in significantly less time at both reduced and simple scavenger community (Fig. 3b) with ‘devil activity’ (HR = 0.92) and wet forest ‘habitat’ (HR = 0.47) suppressing carcass discovery by ravens. Cats showed a similar but weaker response to those observed in ravens (Fig. 3c), however, there were no supported variables.

**Figure 3.**
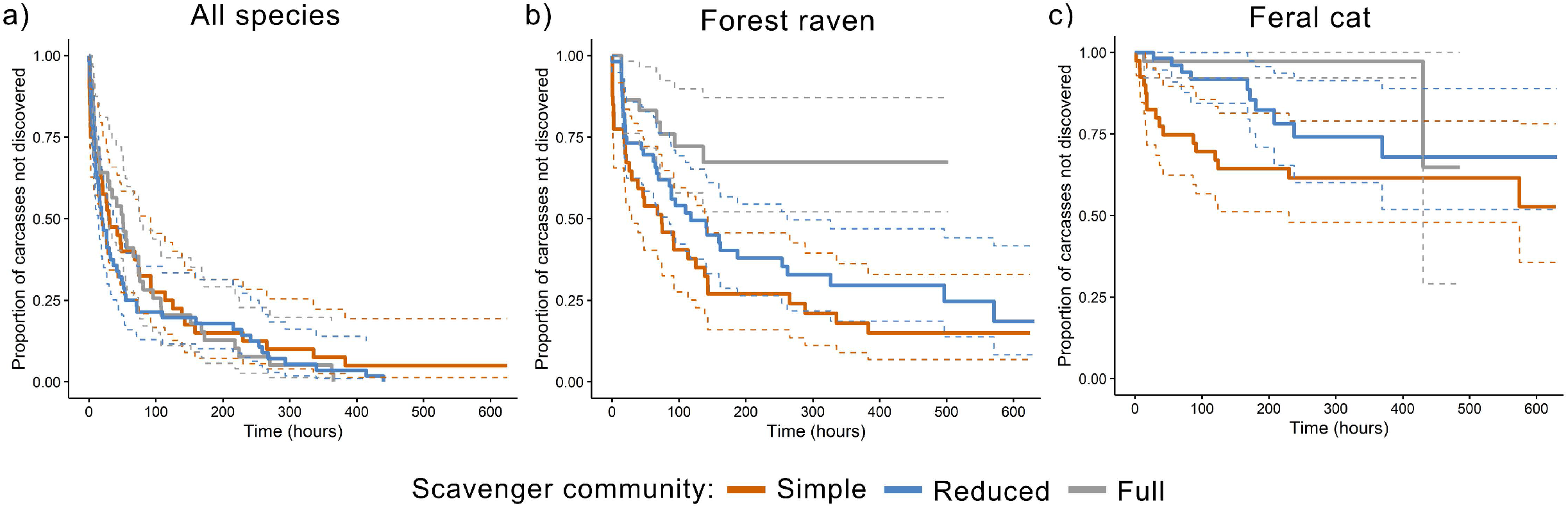
The proportion of carcasses not discovered by (a) all species, (b) forest ravens and (c) feral cats for each level of scavenger diversity and abundance. Colour dashed lines indicate the 95% confidence interval.

### ii) Carcass use

Forest ravens were the main beneficiary of the absence of native mammalian carnivores (Table 1; Fig. 4a). As a proportion of total foraging time for all species, we found that ravens fed almost twice as long in the simple scavenger community (88.2% of total foraging time by all species) compared to ravens in the reduced scavenger community (47.9%) and five times as long as in the full scavenger community (17.3%; Fig. 4d). Both devil scavenging time (Std. coef = 0.53; ES: -0.99; Fig. 4b) and devil activity (Std. coef = 0.47; ES: -0.98; Fig. 4c) had a strong negative effect on the likelihood of a raven feeding at a carcass. However, only devil scavenging time impacted total duration of raven scavenging (Std. coef = 0.81), having an overall negative effect (ES: -0.99; Fig. 4e).

**Table 1.**
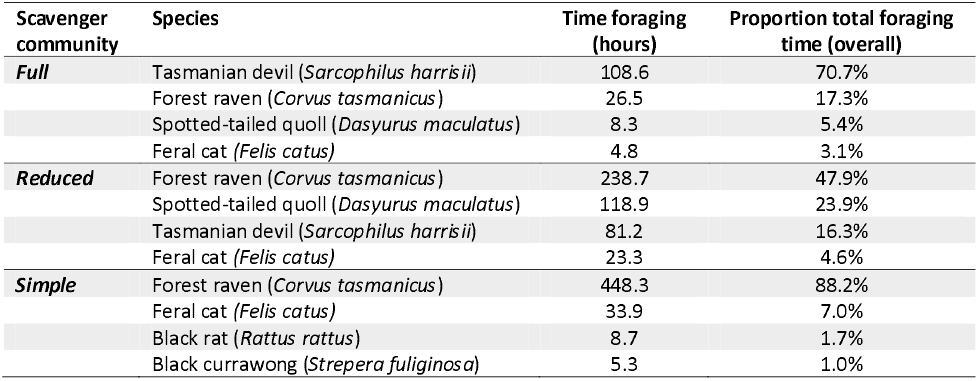
Total foraging times and proportion of total foraging time for the top four scavengers in each scavenger community.

**Figure 4.**
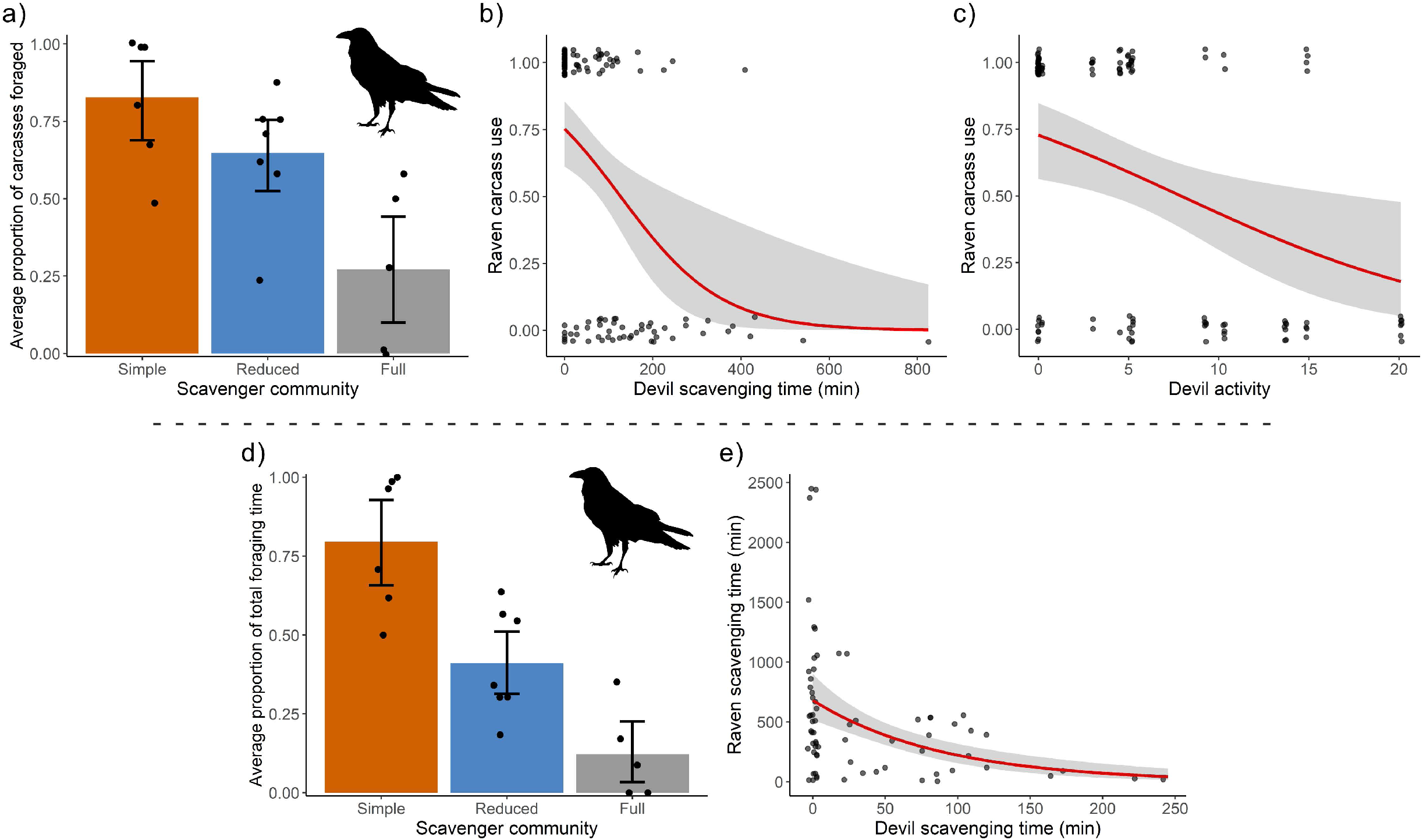
Carcass consumption by forest ravens. Top row: (a) the proportion of total carcasses foraged, with the response curves of the best predictors, being (b) devil scavenging time and (c) devil activity. Bottom row: (d) the average proportion of total foraging time with the response curve of the best predictor, (e) devil scavenging time. In (a) and (d) each dot corresponds to the mean value for the study sites and error bars are bootstrapped 95% confidence intervals.

Feral cats scavenged on a higher proportion of carcasses in the absence of native mammalian carnivores. Cats in the simple scavenger community fed on 44% of carcasses, which was 2.4 times more than in the reduced community region (19%), and eight-fold more than in the full community region (6%; Fig. 5a). The probability of cats scavenging was best predicted by devil activity (Std. coef = 0.65) and the relationship was negative (ES: -0.91; Fig. 5b).

**Figure 5.**
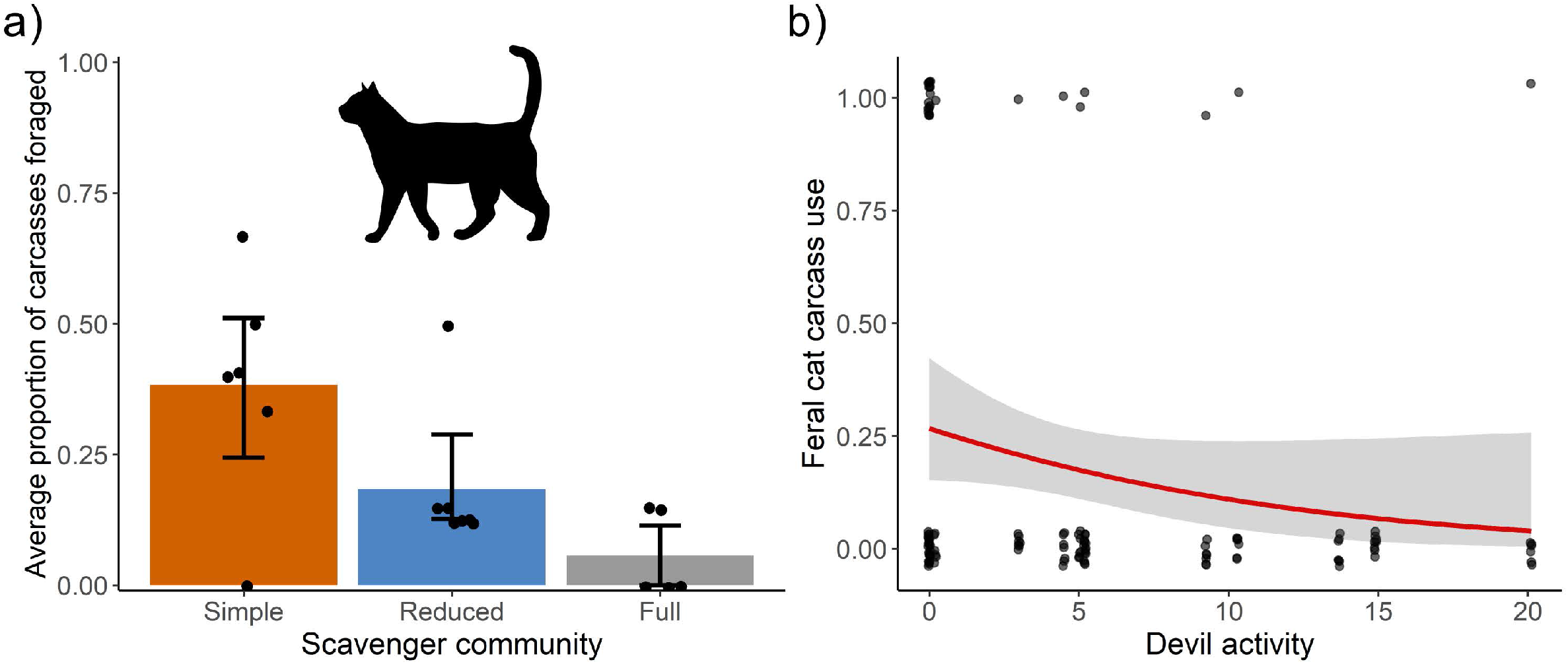
Carcass consumption by feral cats: (a) the proportion of total carcasses foraged, and (b) the response curves of the best predictor, devil activity. In (a) each dot corresponds to the mean value for the study sites and error bars are bootstrapped 95% confidence intervals.

## Discussion

We used a naturally occurring experiment, of reduction or extirpation of native mammalian scavengers, to examine the effects on scavenging by avian and invasive-mammalian scavengers. The apex mammalian scavenger, the Tasmanian devil, had an overwhelmingly dominant effect on scavenger dynamics, with carcasses persisting 4.6 times longer in areas with a simple scavenger community structure. Smaller scavengers, most notably forest ravens, were the main beneficiary of native mammalian carnivore loss. Further, invasive cats scavenged almost 50% of all carcasses in areas with simplified carnivore communities, highlighting that—contrary to general wisdom [36, 37]—scavenging can contribute an important source of food for cats. This also suggests potential avenues for reducing the cat’s devastating effects on native wildlife. Overall, this research highlights the crucial role of scavenging by larger mammals and shows that their effects are not easily replaced once lost. Rewilding of larger carnivores could restore their function within an ecosystem and provide top-down control on mesoscavenger populations (Fielding et al. 2020).

We found that smaller mesoscavengers were unable to replicate the scavenging efficiency of the larger, specialist scavenger, the Tasmanian devil [21]. In the absence of native mammalian carnivores, carcasses persisted for almost five times longer than in areas with higher mammalian carnivore diversity (Fig. 2). While a higher proportion of carcasses in the simple community region were of the larger Bennett’s wallaby, the several Tasmanian pademelon carcasses we used also persisted until the end of the study. Previous research has found that mesoscavengers, including corvids, were unable to functionally replace raptors in urban areas, with 70% of fish carcasses remaining [38]. Additionally, the experimental exclusion of carcasses from vultures resulted in 10 times as many carcasses not fully consumed by the remaining scavengers [39]. While all carcasses within the simple community region in our study persisted until the study’s end, only 5% of carcasses were not discovered and scavenged upon by the remaining mesoscavengers (Fig. 3a). Clearly, the absence of native mammalian carnivores had little impact on the remaining scavengers locating and feeding upon the carcasses, thereby indicating little facilitation of carrion resources by top scavengers through advertising or increased accessibility.

We found evidence that top carnivores limit carrion access for smaller scavengers. Ravens were the main beneficiary of native mammalian carnivore loss, with ravens in the simple community region finding 88% of the carcasses (Fig. 3b) and foraging for seven times longer than in the full scavenger community regions (Fig. 4d). Devils suppressed raven carcass utilisation (Fig. 4e), probably because nocturnal devils consumed the resources before diurnal ravens discovered them. This finding supports previous findings that under low levels of competition raven populations on the Bass Strait islands prioritise scavenging on roadkill across the entire year even when other resources (e.g., invertebrates, fruit, seeds) are available [40].

Until recently, cats were believed to rarely scavenge [36, 37]. However, there is now a growing body of evidence that they actively scavenge, especially when they perceive little risk [41]. Our data show that in the simple scavenger community, feral cats scavenged at eight times the rate of cats living in a full scavenger community (Fig. 5a). Furthermore, the presence of devils in the landscape suppressed cat scavenging behaviour, potentially through interference competition (cats also being mostly nocturnal foragers). Past research has demonstrated that the presence of devils within an environment can trigger avoidance strategies in cats to evade interspecific conflict [14, 42, 43]. Reduced interference competition caused by top-carnivore loss can have cascading effects throughout an ecosystem, potentially leading to population increases of, and expanded functional roles for, smaller carnivores [44]. However, fear effects imposed by larger carnivores on mesocarnivores are not fully understood, and further studies are required to disentangle these dynamics between carnivores [18]. Combining several lines of investigation (e.g., GPS data on multiple predators combined with cameras on carcasses) would help quantify the risk-reward trade-off of carcasses, and could help reveal under what circumstances carcasses are “fatally attractive” to mesopredators [19].

Following the disease-driven decline of devils across Tasmania, quolls also increased their use of carrion in areas of low devil density [14]. Indeed, in areas of greater devil decline, such as north-Eastern Tasmania, quoll scats contained many large-mammal remains, suggesting that the loss of top scavengers improved scavenging opportunities for quolls [45]. Despite quolls being mesoscavengers, they have, like devils, also been documented chasing cats from carcasses, providing evidence of interference competition [14]. However, we only found a weak effect of quoll abundance on carcass use by ravens and cats. As quolls are non-specialised and smaller scavengers, they are much less efficient than devils and it is therefore difficult for them to monopolise a carcass in the same way [46]. Additionally, the effects of devils—as a dominant and specialised scavenger— on other opportunistic scavengers might simply be too strong, acting to mask any potential impacts the quoll may have on cats and ravens [21].

As highly efficient scavengers, the loss of apex scavengers can lead to increased food availability for mesoscavengers which could result in increases in abundance [1]. For example, the absence of vultures (*Gyps* spp.) in south-eastern Spain led to a higher abundance of red foxes (*Vulpes vulpes)* due to greater availability of carrion [47]. Similarly, feral dogs (*Canis lupus*) and rodents have increased in abundance in areas of India due to a rise in carrion availability following widespread vulture declines [48]. In the Bass Strait region, anecdotal evidence suggests that forest-raven and feral-cat populations are growing on King Island [28, 49]. Enhanced opportunities to feed to on roadkill [40] and other carrion, as demonstrated in this study, may provide explanations for this apparent increase in abundance. Further research is needed to confirm whether these species are truly increasing in abundance. Elevated numbers of forest ravens could have destructive effects for the local birds on the islands through heightened levels of depredation, and impact local farmers through increased attacks on livestock, as shown in other corvid studies [41, 50], as well as on King Island specifically [51]. While past research in Tasmania found no impact of forest raven abundance on the abundance of other bird species [52], these impacts may differ on the Bass Strait islands if the raven population size is greater. Meanwhile, the impacts of the invasive feral cat on small mammals or birds are well-documented, with many species now threatened with extinction or already lost due to heightened predation risk [53, 54]. Despite these apparent increases in abundance, feral cats and forests ravens are less efficient scavengers than devils [46], meaning carcasses may persist in an environment for longer. This could have adverse effects on both animal and human health due to the increased spread of carrion-borne diseases [1, 17, 55].

Large carnivore populations have fluctuated due to human persecution and habitat loss, causing trophic cascades throughout food webs across the globe [6]. In our study, we found that top scavengers, like Tasmanian devils, limit carrion use and discovery by smaller scavengers, such as ravens and cats. However, it remains unclear how this may impact mesoscavenger population abundance and whether there are cascading effects on small prey species. In the absence of top mammalian scavengers, we found that carcasses persisted beyond the study length (∼ 3 weeks). Further research is required to see how this may impact the transmission of carrion-borne diseases and scavenging by invertebrates. In this mensurative experiment, we demonstrate that the absence of top mammalian scavengers results in the loss of essential ecosystem functions, providing support for novel management approaches, such as trophic rewilding [56-58]. Overall, our findings further highlight and clarify the integral role native mammalian scavengers perform within an ecosystem, demonstrating the ecological significance of global mammalian carnivore conservation.

## Acknowledgements

The authors would like to acknowledge the palawa peoples of lutruwita, the traditional custodians of the lands on which this work was completed. We extend our gratitude to the many fieldwork volunteers. We would also like to thank the Flinders Island Aboriginal Association Inc. (FIAAI) and various landowners across Tasmania who allowed us to complete fieldwork on their properties. This work was funded by the Holsworth Wildlife Research Endowment and Australian Research Council (ARC) grants FL160100101, CE170100015 to B.W.B. and DP110103069 to M.E.J.

