## Supplementary Information for "Dominant carnivore loss benefits native avian and invasive mammalian scavengers"

**Supplementary Table 1** Model selection table for mixed-effects Cox proportional hazards models. These model sets were used against four different response variables – discovery of carcasses by all species; discovery of carcasses by forest ravens; discovery of carcasses by feral cats and persistence of carcasses. All models included site location as a random effect to account for variation across the study sites.

|  | **Model structure** |
| --- | --- |
| n | null |
| 1 | devil activity |
| 2 | habitat |
| 3 | quoll activity |
| 4 | devil activity + quoll activity |
| 5 | quoll activity + habitat |
| 6 | devil activity + habitat |
| 7 | devil activity + quoll activity + habitat |
| 8 | carcass weight |
| 9 | carcass weight + devil activity + quoll activity + habitat |
| 10 | carcass weight + quoll activity |
| 11 | carcass weight + devil activity |

**Supplementary Table 2** Model selection table for carcass use models. These model sets were used against three different response variables – whether ravens fed at a carcass (GLM with binomial link function); total scavenging time by ravens for the carcasses at which they fed (GLMs with a Gamma distribution and a log link function); and whether feral cats fed at a carcass (GLM with binomial link function). All models included site location as a random effect to account for variation across the study sites.

|  | **Model structure** |
| --- | --- |
| n | null |
| 1 | devil activity |
| 2 | habitat |
| 3 | quoll activity |
| 4 | devil activity + quoll activity |
| 5 | quoll activity + habitat |
| 6 | devil activity + habitat |
| 7 | devil activity + quoll activity + habitat |
| 8 | devil scavenging time |
| 9 | quoll scavenging time |
| 10 | devil scavenging time + quoll scavenging time |
| 11 | devil scavenging time + devil activity |
| 12 | devil scavenging time + carcass weight |
| 13 | quoll scavenging time + carcass weight |
| 14 | devil activity + carcass weight |
| 15 | quoll activity + carcass weight |
